# Genome-AC-GAN: Enhancing Synthetic Genotype Generation through Auxiliary Classification

**DOI:** 10.1101/2024.02.14.580420

**Authors:** Shaked Ahronoviz, Ilan Gronau

**Affiliations:** Efi Arazi School of Computer Science, Reichman University, Herzliya, Israel

## Abstract

In recent years, there have been increasing attempts to develop computational methods for generating synthetic genomic data that aim to mimic real genomic datasets. Artificial genomes (AGs) generated by these methods have emerged as a promising potential solution for privacy concerns raised by public genomic datasets and as means to provide adequate representation of under-sampled populations. However, existing methods for generating AGs provide a very limited capability for faithfully capturing features of different sub-populations within a larger cohort. In this study, we propose a novel method called the Genome Auxiliary Classifier Generative Adversarial Network (Genome-AC-GAN), which generates AGs tailored to specific sub-populations. We conducted experiments to evaluate the performance of the Genome-AC-GAN and compare the AGs it generates with real genomic data as well as with AGs generated by previously published methods. The Genome-AC-GAN outperforms other methods and faithfully models population structure, which is not adequately captured by existing methods. We also demonstrate the use of AGs generated by the Genome-AC-GAN in augmentation of datasets used as training sets for classifying genomes into populations. These experiments demonstrate the benefits of AGs in enhancing classification accuracy, especially when dealing with under-sampled and closely related populations.

## 1 Introduction

The field of genomics has been transformed in the past two decades by the emergence of largescale genomic data sets, such as the 1000 Genomes [37], and the UK Biobank [33]. However, these datasets raise privacy concerns regarding sampled individuals, and they are limited in their capabilities to represent under-sampled populations. In recent years, generative computational models have been suggested as promising solutions for these problems. These models can learn the distribution of genome variants from real largescale datasets and then generate artificial genomes (AGs) that will hopefully capture features of the real distribution. The use of generative models has been shown to be very effective in other domains, such as image synthesis and style transfer [10, 15, 16, 42, 36, 21]. In genomics, the great promise of synthetic data is that if they are sufficiently realistic, they can replace real genomes in downstream analyses. This would potentially reduce the amount of private information leaked from the sampled genomes and allow the generation of sufficient numbers of genomes to alleviate concerns of under-sampling.

Two studies recently published by Yelmen and colleagues examine the utility of different techniques in machine learning in generating AGs [40, 39]. These techniques include generative adversarial networks (GANs) [10], restricted Boltzmann machines (RBMs) [9], and Variational autoencoders (VAE) [18]. The two studies assess the capability of these different approaches in capturing features of real genomes, such as allele frequencies, linkage disequilibrium, and distribution of genomic variants. These comparisons suggests that GANs typically provide the most faithful representation, when compared to real genomes. Another recent study [6] showed that hidden Chow-Liu trees (HCLTs), and their representation as probabilistic circuits, improve the accuracy and efficiency of generating AGs by combining the tractability of a probabilistic model with the expressivity of a deep network. Additional earlier approaches also exist [4, 7, 17], but none of them appear to have significant advantages over the baseline methods considered by Yelmen and colleagues [40, 39].

All existing methods for creating AGs primarily focus on population-level characteristics, and are thus limited in their capabilities to synthesize AGs that capture characteristic features of certain sub-populations within the larger cohort. To address this limitation, we propose here to apply the approach of the auxiliary classifier GAN (AC-GAN), which incorporates in its input a class label that can represent a sub-population of interest. AC-GANs have been shown to be very effective in generating synthetic data for under-sampled populations. For example, a recent study used an AC-GAN to improve the detection of COVID-19 from chest X-rays [38]. An AC-GAN was trained on 721 images taken from healthy individuals and 403 images taken from COVID-19-positive patients. The trained AC-GAN was then used to generate 1399 synthetic images of negative cases and 1669 synthetic images of positive cases. A separate classifier was then trained for COVID-19 detection on the original training set, as well as on a training set augmented with synthetic images generated by the AC-GAN, and the added synthetic images were shown to improve detection accuracy from 85% to 95%. Here, we examine the potential of AC-GANs in improving the representation of under-sampled populations in AGs. We show that the Genome-AC-GAN outperforms existing methods for generating AGs, and we demonstrate its potential for generating AGs that can augment training sets for population classification methods. In particular, we show that the Genome-AC-GAN faithfully captures features of under-sampled populations, which is the main limitation of exiting methods.

## 2 Background

In this section we provide some basic background on GANs, including notations, different variants of GANs, and common training practices.

### 2.1 Standard GAN

A GAN is made up of two separate neural networks: a generator (*G*) and a discriminator (*D*). The generator’s primary objective is to produce synthetic data that closely resembles real data, and the discriminator’s primary objective is to distinguish between real and synthetic data (Figure 1). The generator takes input noise from a simple distribution, *z* ∼ *P*_*noise*_, and transforms it into a data instance, *G*(*z*), which we refer to as an *image*. The discriminator takes in an image, *x*, and determines whether it belongs to the distribution of real images (*x* ∼ *P*_*real*_) or synthetic images (*G*(*z*) | *z* ∼ *P*_*noise*_). The output of the discriminator, *D*(*x*), is the probability that *x* is a real image (and not synthetic). The generator and discriminator are evaluated jointly, according to the following expected log-likelihood function:

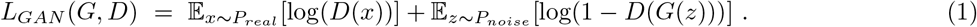

**Figure 1:**
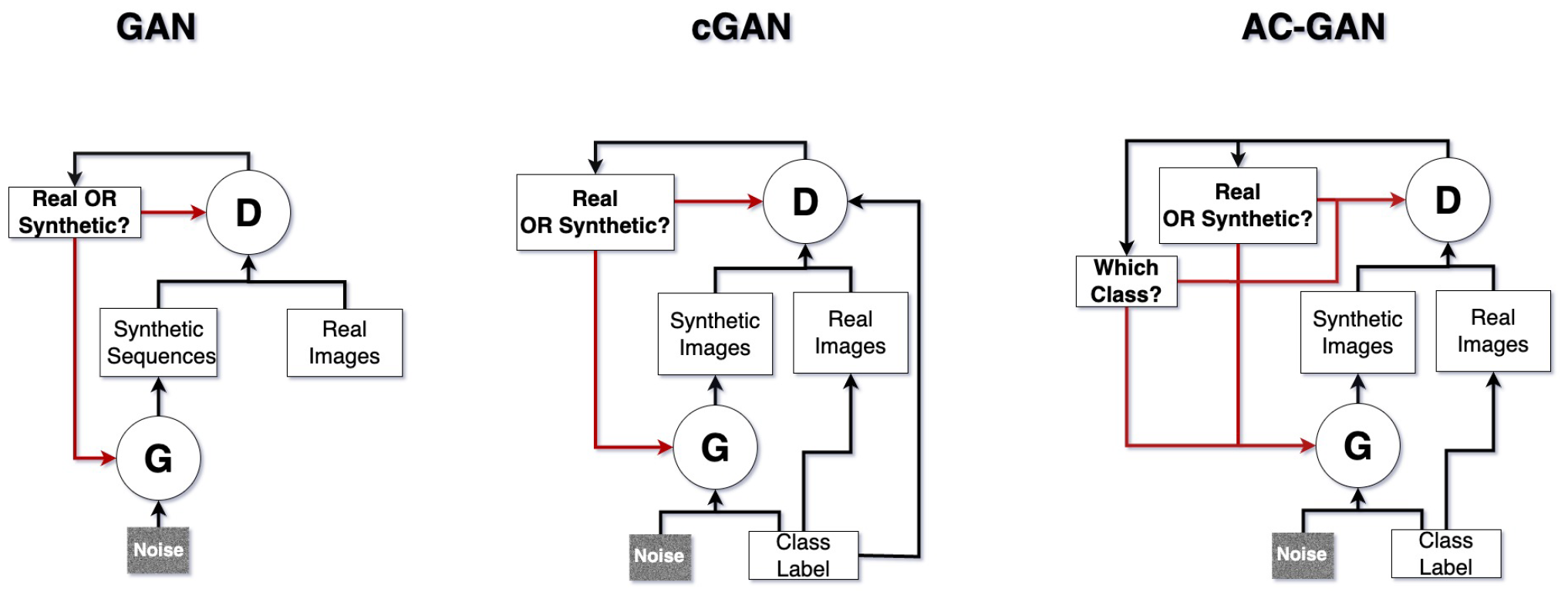
Different types of GANs. In all GANs, the generator (*G*) converts a noise vector into a sequence encoding a synthetic image, while the discriminator (*D*) aims to distinguish synthetic images generated by *G* from real ones. The generator and discriminator are trained simultaneously using feedback regarding the discriminator’s accuracy (red arrows). In conditional GANs (cGANs) and Auxiliary Classifier GANs (AC-GANs), each image is associated with a (discrete) class label. In cGANs, the class is provided as input to both the generator and discriminator, and thus assists the discriminator in its task. In AC-GANs, the class is provided as input only to the generator, and the discriminator is expected to produce it as output. Thus, the discriminator of an AC-GAN is trained to distinguish real images from synthetic ones and also determine the class label of a given image.

The first term in this expected log-likelihood quantifies the expected accuracy of the discriminator in identifying real images, and the second term quantifies the expected accuracy of the discriminator in identifying synthetic images generated by the generator. Note that a good discriminator maximizes the expected log-likelihood, whereas a good generator minimizes it, by making the discriminator wrongly classify synthetic images that it generates as real. Thus, an optimal generator and discriminator may be obtained by solving the following min-max problem:

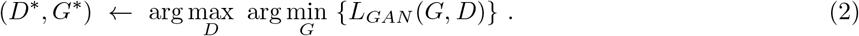

### 2.2 Conditional GAN (cGAN)

A conditional GAN (cGAN) incorporates contextual information during image generation. This approach has been shown to be effective in tasks such as image-to-image translation [13] and attribute-based image synthesis [1]. The contextual information can range from simple discrete class labels to more complex features [25, 28, 26], but for simplicity, we will model it here is a discrete class label, *c*. In cGANs, both the generator and discriminator receive the class label, which consequently improves the realism of the synthetic images generated by *G*, and assists *D* in distinguishing between real and synthetic images. The expected log-likelihood for cGANs is given by considering the conditional distribution of images given each class label, *x* ∼ *P*_*real|*_:

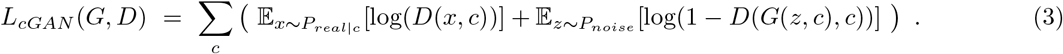

### 2.3 Auxiliary Classifier GAN (AC-GAN)

An auxiliary classifier GAN (AC-GAN) also incorporates class labels as auxiliary information, but unlike in cGANs, the discriminator does not receive the class labels as input, and is expected to produce it as output. Thus, in AC-GANs, the discriminator assumes the responsibility of both discriminating between real and synthetic data and predicting the class label. AC-GANs have been applied in various domains, including image generation for data augmentation and conditional image synthesis tasks [38, 22]. The generator of an AC-GAN is similar to that of cGANs, as it takes a class label and a noise vector as inputs and produces a suitable synthetic image, *G*(*z, c*). The discriminator of an AC-GAN, on the other hand, includes an additional output vector, denoted as *D*_*class*_(*x*), which associates each class label with its predicted probability. Specifically, *D*_*class*=*c*_(*x*) denotes the probability that the discriminator assigns to an input image *x* belonging to class *c*. The expected log-likelihood for AC-GANs considers the joint distribution of images and class labels, (*x, c*) ∼ *P*_*real*_, and is made up of two separate components: a component evaluating the ability of *D* to distinguish synthetic images from real ones (*L*_*distinguish*_), and a component evaluating the classification accuracy of *D* (*L*_*classify*_):

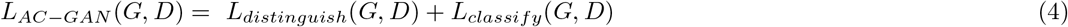

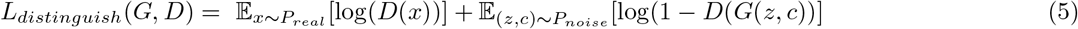

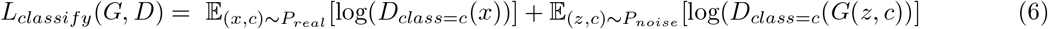

The first component, *L*_*distinguish*_(*G, D*), is similar to that of the standard GAN (see Equation 1). The second component, *L*_*classify*_(*G, D*), measures the accuracy of the discriminator in predicting the class labels associated with the real and synthetic images. The discriminator’s objective is to maximize both components. The generator in an AC-GAN is only adversarial in the distinction task and not the classification task, meaning that its objective is to fool the discriminator to think its generated images are real, but to assist it in correctly predicting the class label. Thus, the objective of the generator is to minimize *L*_*distinguish*_(*G, D*) and maxi-mize *L*_*classify*_(*G, D*). In summary, both the generator and discriminator strive to minimize the classification likelihood in addition to their opposing (min-max) objectives related to discrimination.

### 2.4 Training GANs

Training a GAN involves iterative optimization of the appropriate components of the expected log-likelihood, as presented above. The probability distribution over real images (*P*_*real*_) is represented using a training set. In cGANs and AC-GANs the prevalence of each class in the training set is often set to improve the relative accuracy of generation (and classification) of certain classes of interest. Each iteration of the training procedure involves examining the training set of *n* real images alongside a collection of *n* synthetic images generated by applying the current version of *G* to *n* noise vectors sampled from *P*_*noise*_. The parameters (neural network weights) of the generator and discriminator are modified in each iteration by applying backpropagation to decrease a certain *loss function*. The loss function is designed such that its minimization will increase (or decrease) the appropriate component of the expected log-likelihood. The standard loss function used for GANs, and for the *L*_*distinguish*_ component in AC-GANs, is binary cross entropy (BCE; [11]). In this context, let *y*(*x*) denote the label associated with a given image: *y*(*x*) = 1 if *x* is real and *y*(*x*) = 0 if *x* is synthetic. The contribution of a given image (real or synthetic) to the BCE loss function is defined as follows:

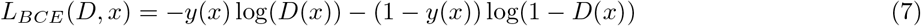

The standard loss function used for the classification task of the discriminator in AC-GANs is the categorical cross-entropy (CCE; [12]), which is a natural extention of BCE to multiple classes. In this context, let 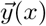 denote the indicator vector for the class label of image 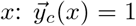 iff the class label of *x* is *c* (and 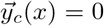 otherwise). Then, if *D*_class_(*x*) denotes the vector of class label probabilities outputted by the discriminator on image *x*, the CCE loss function is defined by the following vector product:

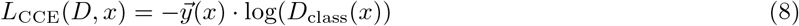

The overall loss considered when updating the discriminator in an AC-GAN is given by summing *L*_BCE_(*D, x*)+ *L*_CCE_(*D, x*) over the *n* training images and *n* synthetic images generated by the current version of *G*. Note that minimizing this loss is expected to increase the expected log-likelihood of Equation 4-6. The overall loss considered when updating the generator in an AC-GAN is given by summing *L*_CCE_(*D, x*) − L_BCE_(*D, x*) over the *n* synthetic images alone. The *n* real images are ignored here because their contribution to the loss is not affected by *G*. The difference in sign stems from the fact that the generator is adversarial to the discriminator in the distinction task (measured by *L*_BCE_) and not in the classification task (measured by *L*_CCE_).

The iterative training procedure is applied until it reaches some convergence. If successful, convergence occurs when the generator produces synthetic images that are indistinguishable from real ones. However, in practice, training GANs can be challenging and time-consuming, and requires careful selection of hyperparameters [3, 19]. One common phenomenon, which is important to avoid is mode collapse, where the generator captures only a subspace of the real image distribution, leading to overfitting and a loss of diversity in the generated images [31].

### 2.5 Polyloss

BCE and CCE are natural candidates for loss functions used when training AC-GANs, because they provide straightforward approximations for the expected log-likelihood in Equations 4-6. However, CCE has a fundamental drawback that can hinder the convergence of the learning process, as illustrated in the following example. Suppose a certain image (*x*) belongs to the second class among four possible classes 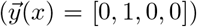. Now consider two different discriminators, *D*^1^ and *D*^2^, with the following outputs:

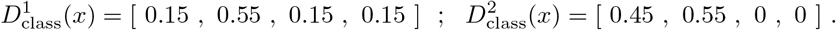

Both discriminators assign the highest probability (0.55) to the true class label and they thus produce the same CCE score: 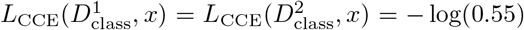. However, the two discriminators assign different probabilities to faulty class labels: *D*^1^ assigns probability 0.15 to each of the three faulty labels, while *D*^2^ assigns a relatively high probability of 0.45 to one of them and probability 0 to the other two. While both discriminators lead to the correct prediction, *D*^1^ produces a more confident prediction, which is something we wish to encourage in the learning process. Thus, we would like to consider loss functions that associate *D*^2^ with a greater loss than that of *D*^1^. The poly-loss cross entropy (PLCE) family of loss functions extends the CCE loss to address this issue by taking into account the probabilities associated with faulty class labels [20]. The PLCE loss is given by adding to the CCE loss a *penalty term* tuned using two hyperparameters, *α* and *ϵ*:

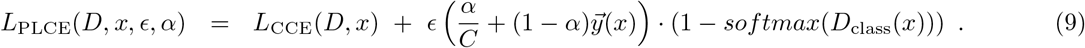

The polyloss penalty is given by a product of two vectors. The first vector is a smoothed version of the class indicator vector 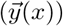, in which all faulty classes are assigned weight 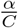 (instead of 0), and the true class is assigned the remaining weight of 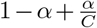 (instead of 1). The second vector is a smoothed version of the output prediction probabilities (*D*_*class*_(*x*)), obtained by applying a softmax transformation and taking the complement to 1. Note that the softmax transformation exponentiates each prediction probability and normalizes the resulting values to sum to 1. Minimizing the polyloss penalty acts to reduce the probabilities assigned to faulty classes, which typically leads to higher confidence in the (correct) classification. Hyperparameter *α* controls the extent to which prediction probabilities for faulty classes influence the penalty term, and hyperparameter *ϵ* controls the contribution of the penalty term to the total loss function. Note that when fixing *ϵ* = 0, we get the standard CCE loss function (regardless of the value of *α*). Selecting effective values for hyperparameters *α* and *ϵ* typically involves various finetuning experiments that examine the convergence properties of the training process.

## 3 Materials and Methods

### 3.1 Genome-AC-GAN architecture

The Genome-AC-GAN model is an adaptation of the basic AC-GAN architecture presented in [25] customized for genotype data. We model genetic data as a sequence of alleles corresponding to a collection of *M* prespecified genomic locations, which will typically be biallelic SNPs. Thus, an individual genome is specified as a binary sequence of length *M*. The architecture of the generator and discriminator is specified below and also illustrated in Figure 2. The input of the generator is a binary noise vector of some pre-specified length (which we set to 600 in all experiments here) and a binary indicator vector (of length *C*) for the class label. The first three (hidden) layers of the generator are of increasing lengths, *M*/1.3, *M*/1.2, and *M*/1.1, and they all utilize the LeakyReLU activation function with an *α* slope of 0.2. The fourth (output) layer uses a tanh activation function and produces a vector of length *M* of values in the rage [− 1, 1]. In the trained generator, this output is converted to binary genetic data by rounding up negative values to 0 and positive values to 1. The discriminator takes in as input a binary vector of length *M* representing a genome, and passes it through three layers of decreasing lengths, *M*/2, *M*/3, and *M*/4. All three layers employ the LeakyReLU activation function with an *α* slope of 0.2. There is a dropout layer after each hidden layer with a dropout rate of 0.2 for the first and second layers, and a dropout rate of 0.1 for the third layer. The discriminator’s validity output (probability of being ‘real’) is provided by applying a sigmoid transformation to the output of the final layer, and the classification output (class probabilities) is provided by applying a softmax transformation. The `regularizers` parameter is set to 0.0001 through the network, and all layers have a pre-specified width (*B*), which corresponds to the batch size used when training the network.

**Figure 2:**
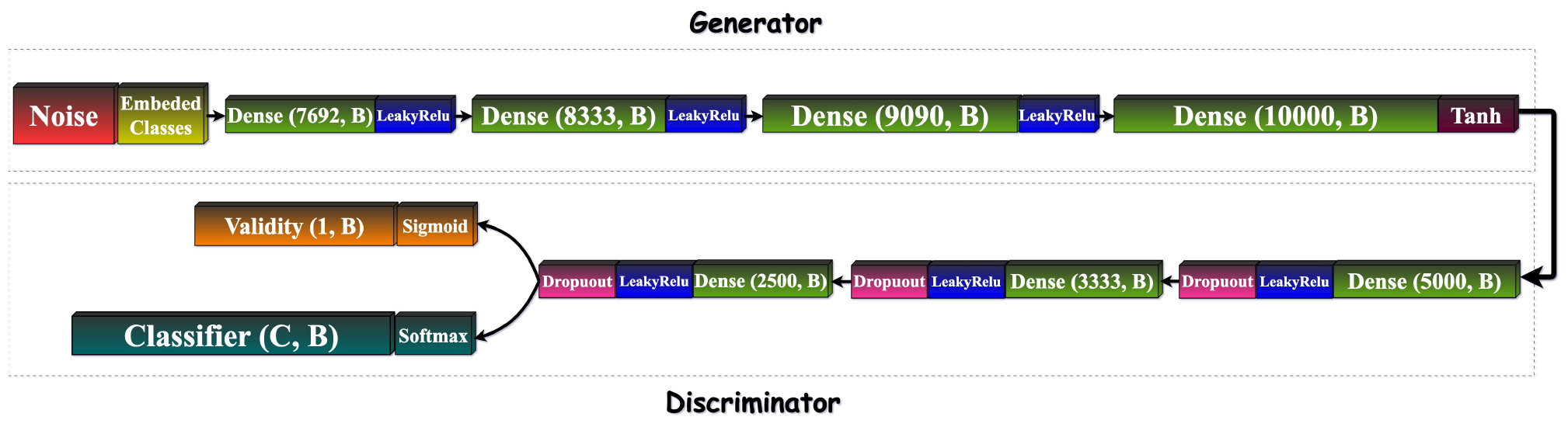
Genome-AC-GAN Architecture. The generator and discriminator of the AC-GAN consist of a sequence of fully-connected neural networks layers. For each layer, we specify its activation function and length. Lengths specified here assume genotype sequences of length *M* = 10, 000, and an arbitrary number of classes (*C*). Each layer also has a width (*B*) corresponding to batch size used during training.

### 3.2 Training procedure

The training process of Genome-AC-GAN adheres to standard practices for AC-GANs with certain modifications. The amount of data considered in each training epoch is determined by the batch size, *B*, which was set to 256 in all experiments reported here. Each epoch involves three stages. First, the loss computed on a set of *B* real genomes from the training set is used to update the parameters of the discriminator. Then, the parameters of the discriminator are updated again using a loss computed on *B* synthetic genomes created by the generator. Finally, a separate collection of *B* synthetic genomes is used to compute a loss for the parameter update of the generator. The BCE and PLCE loss functions are used as described in Sections 2.4 and 2.5, and parameter updates are done using RMSprop optimization. To improve training stability and avoid overfitting, label smoothing is applied, where random noise sampled uniformly in [0, 0.1] is added to the data labels (*y*(*x*) and 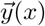) when computing the loss functions [34]. The number of training epochs is set based on visual inspection of convergence of the loss function.

### 3.3 Training and test set

For training and testing, we used data from the 1000 Genomes Project [5], which provides public genome sequences for 2,504 individuals from 26 different national populations worldwide. One individual was assigned to two populations and was removed for our purposes. Each of the remaining 2,503 individuals contributed two separate haploid genome sequences, which we could use for training or testing. Each national population is represented by 120 - 230 haploid genomes, and each of the five continental groups is represented by 690 - 1330 haploid genomes (see Table 1). Following previous studies of methods for generating AGs [40, 39], we focused on 10, 000 single nucleotide polymorphisms (SNPs) within 3 Mbp on chromosome 6 that contain the human leukocyte antigen (HLA) genes. SNPs in this dataset are all biallelic, so each genome (real or synthetic) is represented by a binary sequence of length *M* = 10, 000. To ensure consistency in our experimental comparisons with methods described in [40, 39], we used the exact same phased genome sequences used in these studies, without any additional modifications. We set aside 4,004 genomes for training (∼ 80%) and 1,002 for testing (∼ 20%), making sure that each continental group is split using similar ratios between the two sets. We trained two types of Genome-AC-GAN: one in which class labels correspond to the national populations (*C* = 26 classes), and one in which class labels correspond to continental groups (*C* = 5 classes).

**Table 1:** Overview of the 1000 Genome dataset used for training and testing. The number of haploid genome sequences is specified for each of the 26 national populations, with populations grouped by continent: AFR–Africa, AMR–America, EAS–East Asia, SAS–South Asia, EUR–Europe.

| AFR |  | AMR |  | EAS |  | SAS |  | EUR |  |
| --- | --- | --- | --- | --- | --- | --- | --- | --- | --- |
| pop. | #samples | pop. | #samples | pop. | #samples | pop. | #samples | pop. | #samples |
| GWD | 226 | PUR | 208 | CDX | 186 | ITU | 204 | CEU | 198 |
| ACB | 192 | CLM | 188 | KHV | 198 | PJL | 192 | FIN | 198 |
| ESN | 198 | MXL | 128 | CHB | 206 | STU | 204 | GBR | 182 |
| MSL | 170 | PEL | 170 | CHS | 210 | BEB | 172 | IBS | 212 |
| YRI | 216 |  |  | JPT | 208 | GIH | 206 | TSI | 214 |
| LWK | 198 |  |  |  |  |  |  |  |  |
| ASW | 122 |  |  |  |  |  |  |  |  |
| total | 1322 | total | 694 | total | 1008 | total | 978 | total | 1004 |

### 3.4 Code availability

The Genome-AC-GAN is implemented in Python using the TensorFlow framework, with code accessible via GitHub at: https://github.com/Shaked35/Genome-AC-GAN. The repository contains a complete implementation of the network as well as code required for training the Genome-AC-GAN and reproducing the experiments described in the following section.

## 4 Results

### 4.1 Finetuning the Polyloss penalty

We started with a series of exploratory analyses aimed at finetuning the polyloss penalty in the PLCE loss function (see Section 2.5). In this initial exploration, we found that setting hyperparameters to values *ϵ* = 0.2, *α* = 0.1 leads to effective and efficient training. To demonstrate the influence of the polyloss penalty on the trained AC-GAN, we compare Genome-AC-GANs trained using a PLCE loss function with *ϵ* = 0.2, *α* = 0.1, with a standard CCE loss function (corresponding to setting *ϵ* = 0). In particular, we trained Genome-AC-GANs with *C* = 5 continental population labels for 5, 000 epochs, and examined the classification accuracy of the discriminator.

When comparing the two different Genome-AC-GANs, we see that the model trained with a polyloss penalty attained total classification accuracy 0.88, whereas the model trained without it attained total accuracy 0.82 (Figure 3). The improved accuracy was observed for four of the five continental populations (genomes belonging to EUR were less accurately classified). We wished to confirm that this improved accuracy is observed consistently through the training process and across independent training trials. We thus trained three models using each of the two loss functions, and measured total classification accuracy of the discriminator every 50 epochs (for a total of 5, 000 epochs). Through this process, we obtained for each loss function a total of 303 accuracy measurements. We then removed the top and bottom 2.5% values in each set, and examined the distribution of the remaining 289 values per loss function (Figure 4). A two-sample t-test indicates that the PLCE loss function with hyperparameters *ϵ* = 0.2, *α* = 0.1 leads to significantly higher accuracy throughout training (*p* = 0.00042). Since we posit that higher classification accuracy throughout the training process implies an overall better AC-GAN [14, 31], we use the PLCE loss function with *ϵ* = 0.2, *α* = 0.1 in all Genome-AC-GANs mentioned in the following experiments.

**Figure 3:**
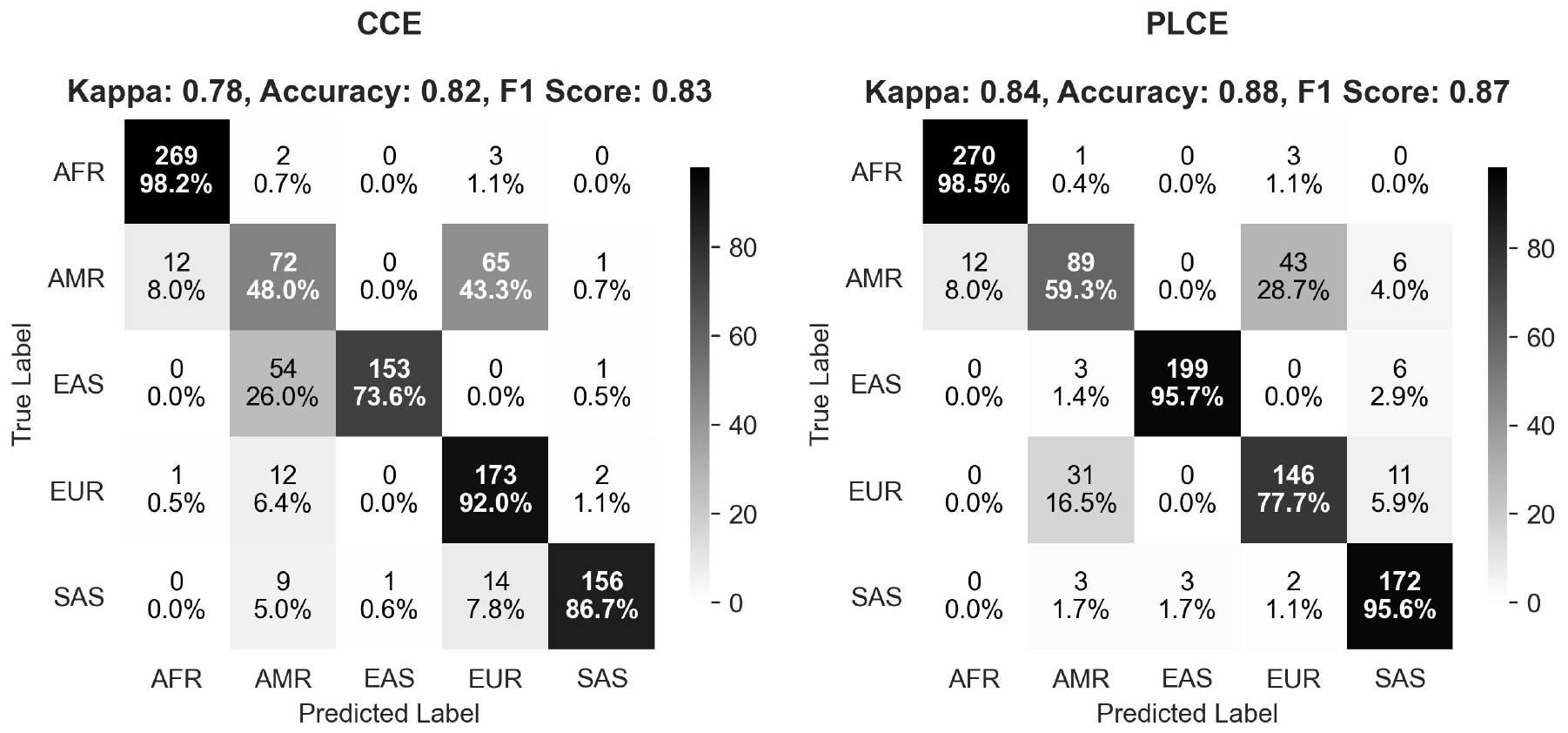
Classification accuracy of Genome-AC-GANs trained using two different loss functions. A Genome-AC-GAN trained using a standard CCE loss function (left) is compared against a Genome-AC-GAN trained using a PLCE loss function with hyperparameters *ϵ* = 0.2, *α* = 0.1. Both models were trained for 5, 000 epochs with *C* = 5 continental population labels. For each Genome-AC-GAN, we report a confusion matrix measuring the classification accuracy of its discriminator on the test set. Rows correspond to the true class label of the test samples and columns correspond to predicted class label (i.e., the class with maximum predicted probability). For each pair of population labels, (*p, q*), we report the absolute number (and percentage) of test samples from population *p* that are predicted to be in population *q*. Above each confusion matrix, we also report the total accuracy, Cohen’s Kappa, and the F1 score [32].

**Figure 4:**
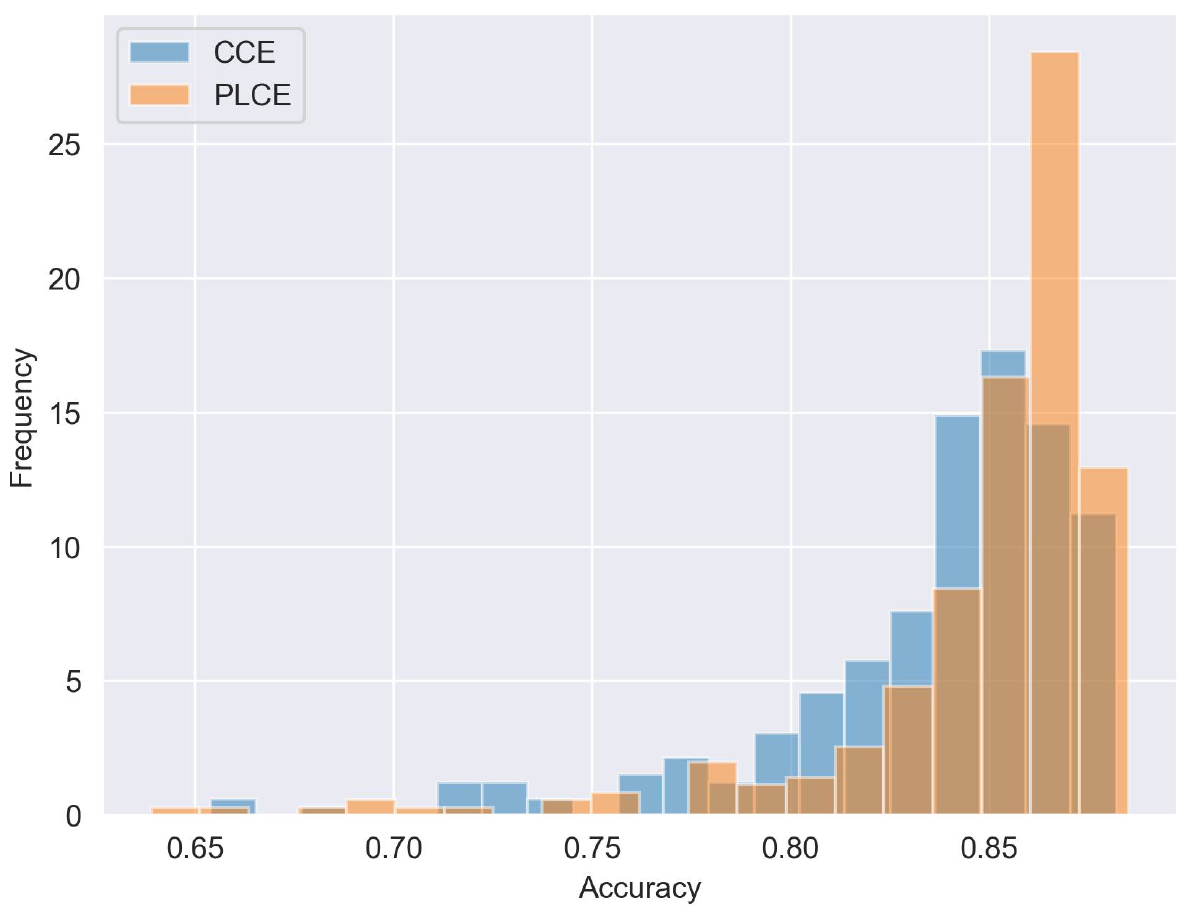
Consistently improved accuracy when training with polyloss penalty. Three Genome-AC-GANs were trained using a standard CCE loss function and three Genome-AC-GANs were trained using a PLCE loss function with hyperparameters *ϵ* = 0.2, *α* = 0.1. For each model, total classification accuracy was measured every 50 epochs for a total of 5, 000 epochs. Measurements were then grouped across the three models trained using the same loss function, and outlier measurements were removed (see text). The distribution of the remaining 289 values is shown here for the standard CCE loss function (blue) and for the PLCE loss function (orange).

### 4.2 Detailed evaluation of artificial genomes generated by the Genome-AC-GAN

We carried out a series of analyses to assess the artificial genomes (AGs) generated by the Genome-AC-GAN. To this end, we trained two different Genome-AC-GANs: One used *C* = 5 continental populations as class labels (AC-GAN-Con), the other used *C* = 26 national populations as class labels (AC-GAN-Nat). Both Genome-AC-GANs were trained using the PLCE loss function with *ϵ* = 0.2, *α* = 0.1 for 10, 000 epochs. Each trained AC-GAN was used to generate 4, 004 AGs, matching the number of genomes and the distribution of class labels in the training set (see Table 1). The two collections of AGs were compared to the collection of real genomes in the test set using a series of measures and analyses that are commonly applied to population genetic data. For reference, we also examined collections of AGs generated by the three top generative models highlighted by Yelmen and colleagues in [40, 39] (Table 2). One of these models (GAN19) was retrained using the exact same training set we used for the two AC-GANs, while for the other two models (RBM23 and WGAN23) we used AGs downloaded from from the authors’ repository. The models that generated these AGs were trained on the same 1000 Genomes data we used, but since we do not know which genomes were used for training these models, there is likely considerable overlap between their training sets and our test set. We note that this gives these two models a slight potential advantage in the comparisons we conducted against the test set. Finally, we also compared the 4, 004 real genomes in the training set to the test set, to provide an upper bound on the expected similarity, since AGs are not expected to appear more realistic than real genomes on which they were trained.

**Table 2:** Artificial genomes used in comparative analysis. AC-GAN-Con and AC-GAN-Nat are the two AC-GANs we developed, trained on the same training set, but with different class label resolutions. Other models were highlighted in recent studies [40, 39] for their robust performance. The GAN19 model was retrained on the same training set as our AC-GANs, while AGs for the RBM23 and WGAN23 models were downloaded from the code repository of [39]. All five sets of AGs were compared using a series of measures and analyses against the test set as well as the training set.

| Model | Description |
| --- | --- |
| AC-GAN-Con | An AC-GAN trained with $C = 5$ continental groups as class labels. 4,004 AGs were generated using this model |
| AC-GAN-Nat | An AC-GAN trained with $C = 26$ national populations as class labels. 4,004 AGs were generated using this model |
| GAN19 | A GAN based on the architecture presented in [40], which was re-trained using the process described in [40] on the same training set as the two AC-GANs. 2,048 AGs were generated using this model |
| RBM23 | A Restricted Boltzmann Machines (RBMs) described in [39]. An RBM is a probabilistic graphical model with two layers of random variables: a visible layer representing observed variables (the genotypes in our case) and a hidden layer representing latent variables (a noise vector). The two layers are typically fully connected. The model was not retrained. Instead, we used AGs generated by a pretrained RBM and published by Yelmen and colleagues. The RBM was trained on the same 1000 Genomes data we used for training and testing, but the actual set used for training was different from the one we used for the AC-GANs and thus likely overlaps with our test set. 5,000 AGs were downloaded for this model from https://gitlab.inria.fr/ml_genetics/public/artificial_genomes/-/blob/master/RBM_AGs/10K_SNP_RBM_AG_1050epochs.hapt.zip |
| WGAN23 | A Wasserstein GAN (WGAN) described in [39]. WGAN uses a loss function which considers the Wasserstein distance between real and generated data. This approach has been shown to improve robustness and ease of training. As with RBM23, the model was not retrained. Instead, we used AGs generated by a pretrained WGAN and published by Yelmen and colleagues. The WGAN was trained on the same 1000 Genomes data we used for training and testing, but the actual set used for training was different from the one we used for the AC-GANs and thus likely overlaps with our test set. 5,008 AGs were downloaded for this model from https://gitlab.inria.fr/ml_genetics/public/artificial_genomes/-/blob/master/GAN_AGs/10K_SNP_WGAN_AG.hapt.zip |

#### Genetic variation by principal component analysis

We started by examining the overall genetic variation as depicted by principal component analysis (PCA). We wished to see how well different collections of AGs captured the typical pattern of variation observed when projecting population genetic data onto its two major principal components (PCs) [24]. From the initial sets of AGs generated by each model (Table 2), we resampled 50 subsets of 1, 002 AGs, matching the size of the test set. Each set of 1, 002 AGs was then projected onto its two main PCs and compared to the projection of the real genomes in the test set by computing the Wasserstein distance between them [27]. For comparison purposes, we did the same resampling process with the genomes in the training set. The distribution of Wasserstein distances for each generative model is shown in Figure 5. We observe that the Wasserstein distances of AGs generated by the two AC-GANs (906 ± 62 and 920 ± 79) are only slightly larger than those of the training set (897 ± 69). The distances observed for GAN19 and RBM23 are significantly larger (2662 ± 43 and 1657 ± 166), indicating a considerably poorer fit. For the WGAN23 model, the Wasserstein distance of an average collection of AGs is only slightly larger than that obtained by our AC-GANs, but the variation is larger (992 ± 123). This observation highlights the need to examine the variance of features of AG sets and not just their expected values. Interestingly, the WGAN23 model did not outperform our AC-GANs despite the fact that its training set likely had a significant overlap with our test set. Thus, the genetic variation represented by the test set is best captured by the two AC-GANs, which shows only a slight degradation relative to the real genomes in the training set. This is illustrated in Figure 6, where we plot for each model the set of resampled AGs with the smallest Wasserstein distance from the test set. Here, we see that the two AC-GANs appear to better capture some of the subtleties in the distribution of real genomes, such as the gap between the cluster representing African populations (bottom right of PCA) and other populations.

**Figure 5:**
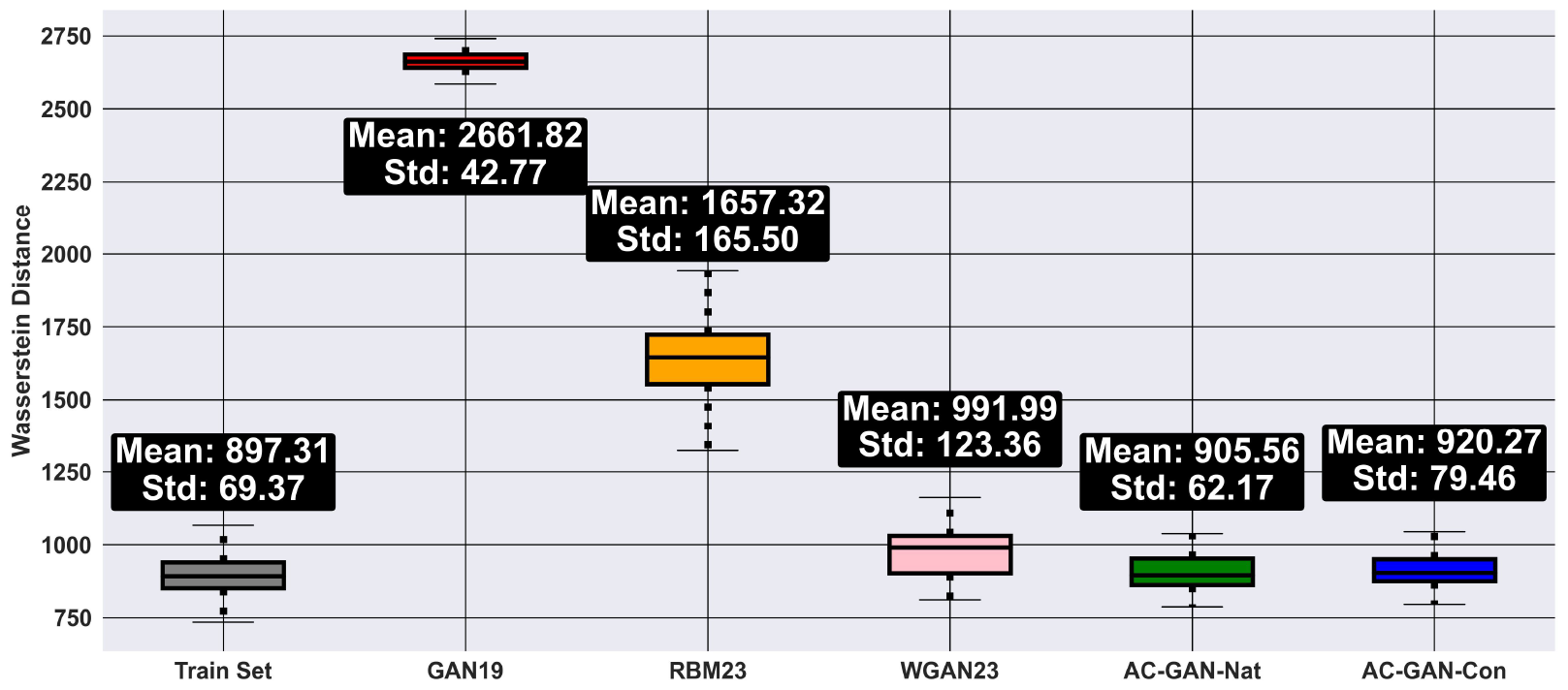
Wasserstein distance between PCA projections of AGs and real genomes. For each collection of AGs, 50 subsets of size 1, 002 were resampled and projected onto their two main PCs together with the test set. The boxplots depict the median, quartiles and outliers of the Wasserstein distances between the two projections across the 50 resampled sets, with the mean and standard deviation specified above each boxplot. A similar analysis was conducted with the training set to provide an estimate for best expected performance.

**Figure 6:**
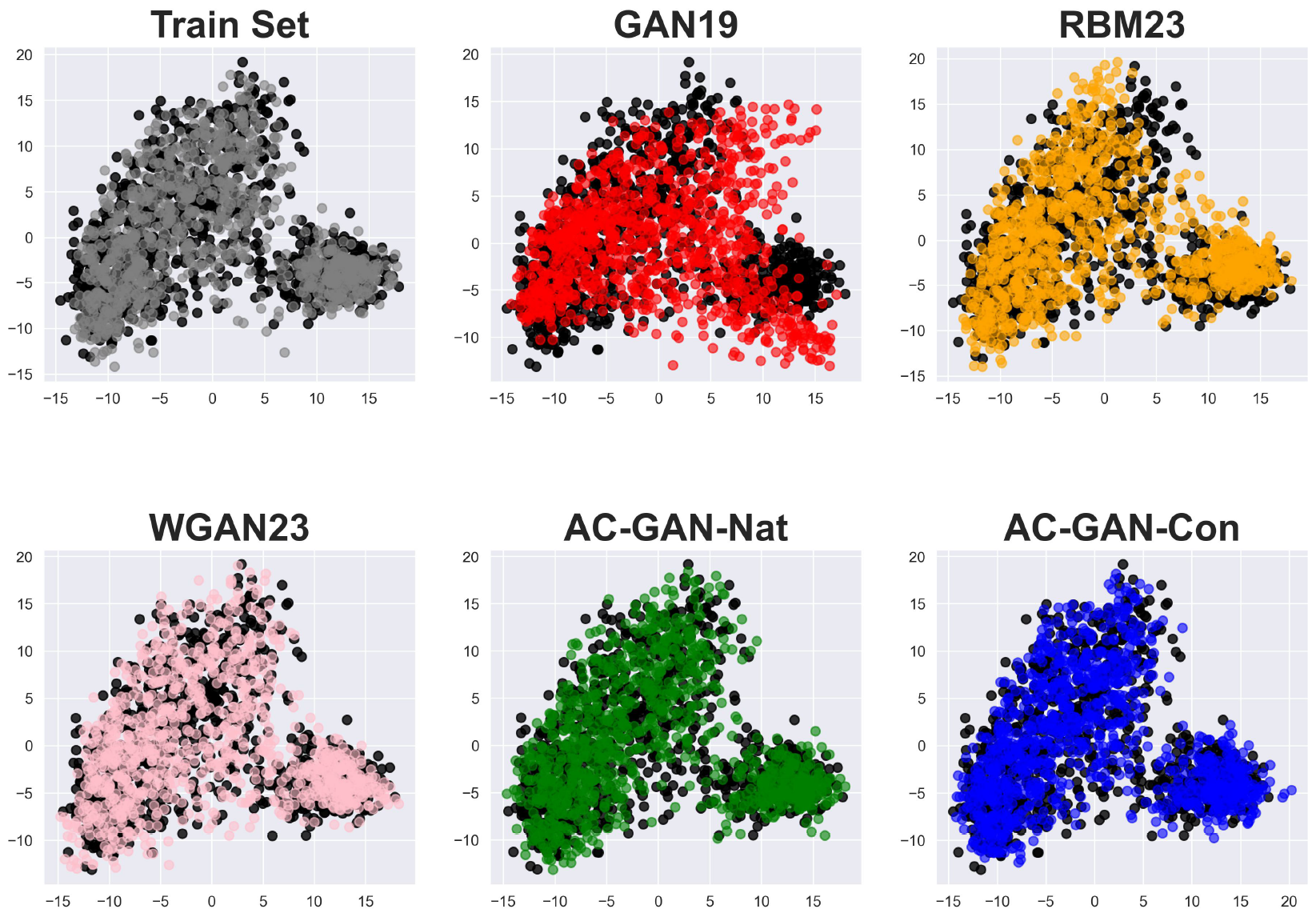
PCA for optimal set of AGs per generative model. Each collection of 1, 002 AGs was projected onto its two main PCs together with the 1, 002 genomes in the test set (black circles in the background). For each generative model, we show the collection of AGs (out of 50) with minimum Wasserstein distance from the test set (see Figure 5).

Unlike previous models for generating AGs, our two Genome-AC-GANs explicitly model genetic variation within and between continental groups. We examined how well this is done by conducting additional PCAs for all real genomes and AGs generated by AC-GAN-Con and AC-GAN-Nat, and coloring each genome based on its continental group (Figure 7). For this purpose, we merged the test and training set to view all 5, 006 real genomes, and used all 4, 004 AGs for each AC-GAN. In AC-GAN-Con, we colored each AG according to the continental group associated with its class label, and in AC-GAN-Nat, we colored each AG according to the continental group to which its national population belongs (see Table 1). The distribution of the different continental groups in the two sets of AGs closely follows the distribution observed in the real genomes and is consistent with previously reported spatial distribution [24]. This observation suggests that genetic variation within and between different continental groups is faithfully modeled by the two Genome-AC-GANs using the national and continental class labels. For a more detailed comparison, we examined genetic variation for each population separately (Figure 8). For each of the five continental groups, we projected AGs generated by AC-GAN-Con under that class label onto the main two PCs, together with real genomes from the test and training set from that continental group. We then did the same thing with AGs generated by AC-GAN-Nat under each of the 26 national populations. Real genomes from the training set were included in this comparison because some of the national populations were sparsely represented in the test set. A visual comparison of AGs from AC-GAN-Con and real genomes confirms that the genetic variation of each continental group is captured with reasonable accuracy. The performance of AC-GAN-Nat appears to vary across national populations. For most populations, the real genetic variation is captured quite well by the AGs. This is particularly impressive in the case of the populations that have a multi-modal distribution in the 2D PCA, such as PJL, KHV, and BEB. However, the AGs generated for some populations with multi-modal distributions, such as PUR, ACB, and ASW, appear to be modeling well only the main modes of the distribution, at the expense of the secondary modes. This is perhaps not very surprising, given the fact that the training set typically includes fewer than 200 genomes from each of these populations (roughly 80% of the genomes in each population; see in Table 1). Thus, our analysis suggests that the Genome-AC-GAN is effective in modeling genetic variation within and between multiple classes, as long as classes are represented reasonably well in the training set and do not have an overly complex pattern of genetic variation.

**Figure 7:**
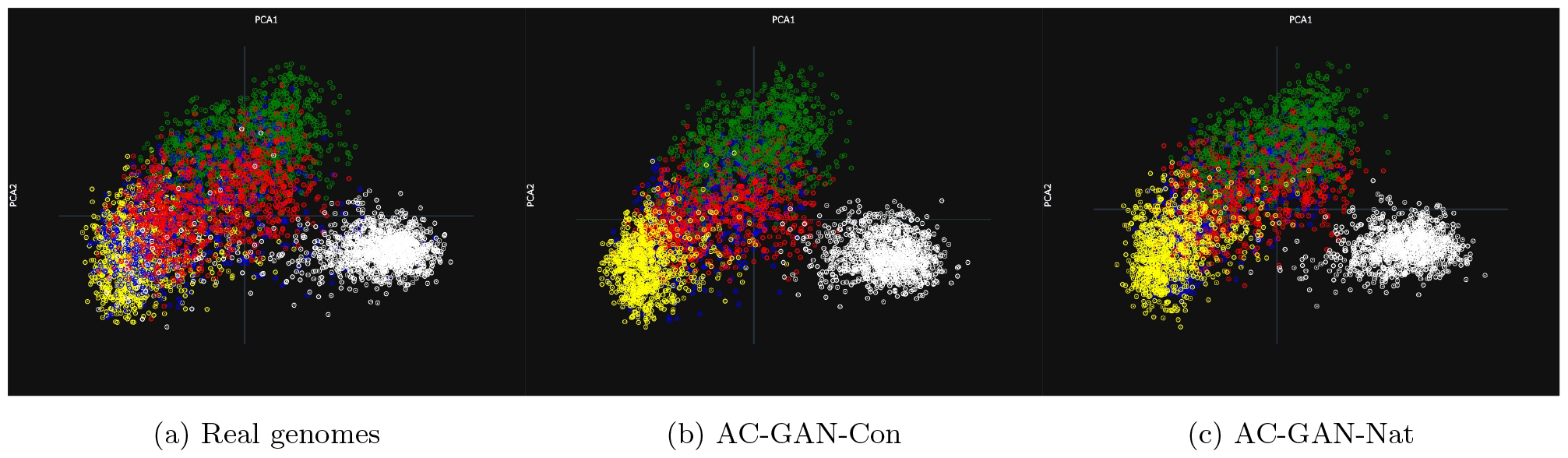
PCA with samples labeled by continental group. Sets of real and synthetic genomes projected onto their two main PCs and colored by continental group: AFR – white, AMR – blue, EAS – green, EUR – yellow, SAS – red. (a) 5, 006 real genomes from the training and test sets. (b) 4, 004 AGs generated by AC-GAN-Con. (b) 4, 004 AGs generated by AC-GAN-Nat.

**Figure 8:**
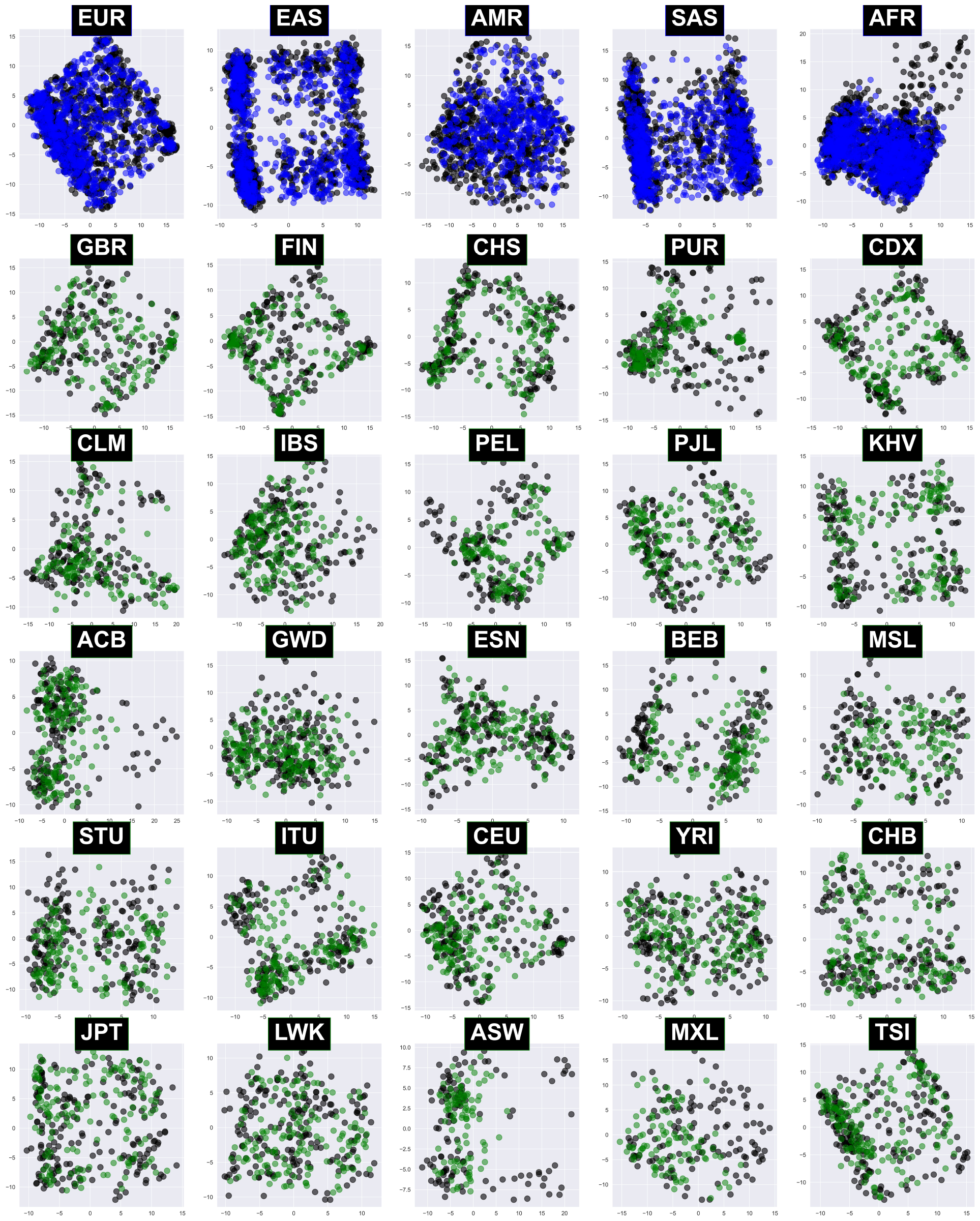
PCA by population. AGs generated for a specific population class label were projected onto two main PCs together with all genomes from the training and test set with the same population label (black circles in the background). The top five panels show AGs generated by AC-GAN-Con (blue) partitioned according to each of the five continental groups. The remaining 25 panels show AGs generated by AC-GAN-Nat (green) partitioned according to 25 national populations. The GIH population from South Asia (Gujarati Indians in Houston) was omitted from this visualization to obtain a 5 *×* 5 layout. It produced very similar distributions for its AGs and real genomes as ITU (Indian Telugu in the U.K.).

#### Allele frequencies

After confirming that the Genome-AC-GAN adequately models global patterns of genetic variation, we wanted to evaluate its effectiveness in capturing the allele frequencies in each of the 10, 000 genomic locations. This is a highly desired feature for AGs, especially for downstream analyses that concern low-frequency alleles [35]. Thus, for each of the 10, 000 SNPs, an arbitrary allele out of the two was selected and its frequency was measured in the test set. These (real) allele frequencies were compared against the frequencies measured using each of the five collections of AGs, and frequencies measured using the training set (Figure 9). Overall, all models for generating AGs appear to produce allele frequencies highly correlated with real ones, with correlations above 98% for all methods. The deviations from real genomes in all models is most prominent in SNPs with allele frequencies above 0.8 and below 0.2. When focusing on these SNPs, we see that the WGAN23 model produces the most accurate allele frequencies. The two AC-GANs produce slightly more noisy allele frequencies, with AC-GAN-Nat producing slightly more accurate allele frequencies when compared to the AC-GAN-Con, likely due to its higher resolution classes. The other two generative models (GAN19 and RBM23) produce somewhat less accurate allele frequencies. As mentioned previously by Yelmen and colleagues in [39], the noise in allele frequencies in all models (with the possible exception of WGAN23) is biased toward the extremes: low-frequency alleles are under-sampled, and high-frequency alleles are over-sampled. Thus, Genome-AC-GAN provides an adequate model for allele frequency, but shares some of the limitations of previously published generative models.

**Figure 9:**
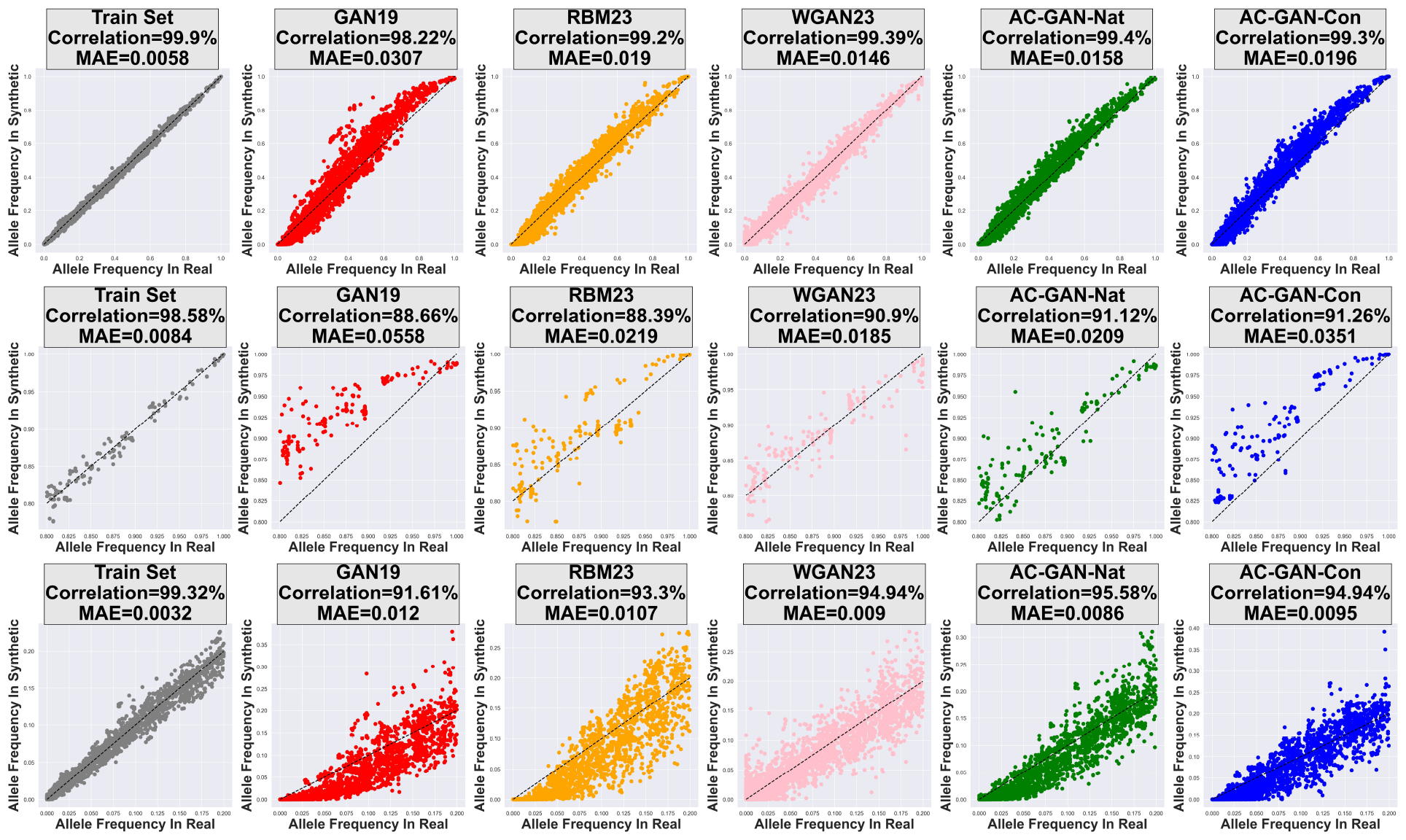
Allele frequencies in real and synthetic genomes. The allele frequency in each of the 10, 000 SNPs was measured for AGs generated by each of the five methods (Table 2) and plotted against the allele frequency measured for that SNP in the test set. In each SNP, one of the two alleles was arbitrarily selected to be measured here. In each plot, the X axis corresponds to the frequency in the test set, the Y axis corresponds to the frequency in the set of AGs, and the correlation and the mean absolute error (MAE) between the two frequencies are reported. Allele frequencies were also computed for the training set to provide an estimate for best expected performance. The top row shows all SNPs, the middle row shows SNPs with real allele frequencies greater than 0.8, and the bottom row shows SNPs with real allele frequencies smaller than 0.2.

A unique feature of our AC-GANs, when compared to other generative models, is their ability to model population-specific allele frequencies [23]. To evaluate how well this is achieved by our AC-GANs, allele frequencies were measured for each SNP in the five continental groups and the 26 national populations (Figure 10). Since national populations are sparsely represented in the 1000 Genomes data (see Table 1), expected allele frequencies were computed using all 5, 006 real genomes in the test and training set. The continent-specific allele frequencies measured in the AGs generated by AC-GAN-Con were highly correlated with those measured in real genomes (> 98%) in all five continental groups, similar to what we saw globally in Figure 9. Producing accurate allele frequencies for national populations, which are more sparsely sampled, appeared to pose a bigger challenge to AC-GAN-Nat. Correlations were above 98% for 12 of the 26 national populations, and above 96% for all populations other than PUR (Puerto Rico), for which the correlation was 94%. The reduced accuracy for this population possibly stems from the fact that it experienced recent admixture, making it more difficult to characterize. Overall, we observe that the AC-GAN model extends the capability of previous generative models to model allele frequencies to population-specific allele frequencies. Accuracy of allele frequencies appears to be higher when using more coarse-grained classes (as in AC-GAN-Con), but it remains reasonably accurate (correlation above 96%) even for classes with fewer than 200 training samples, such as MXL (Mexican ancestry in Los Angeles) and ASW (African Ancestry in Southwestern USA).

**Figure 10:**
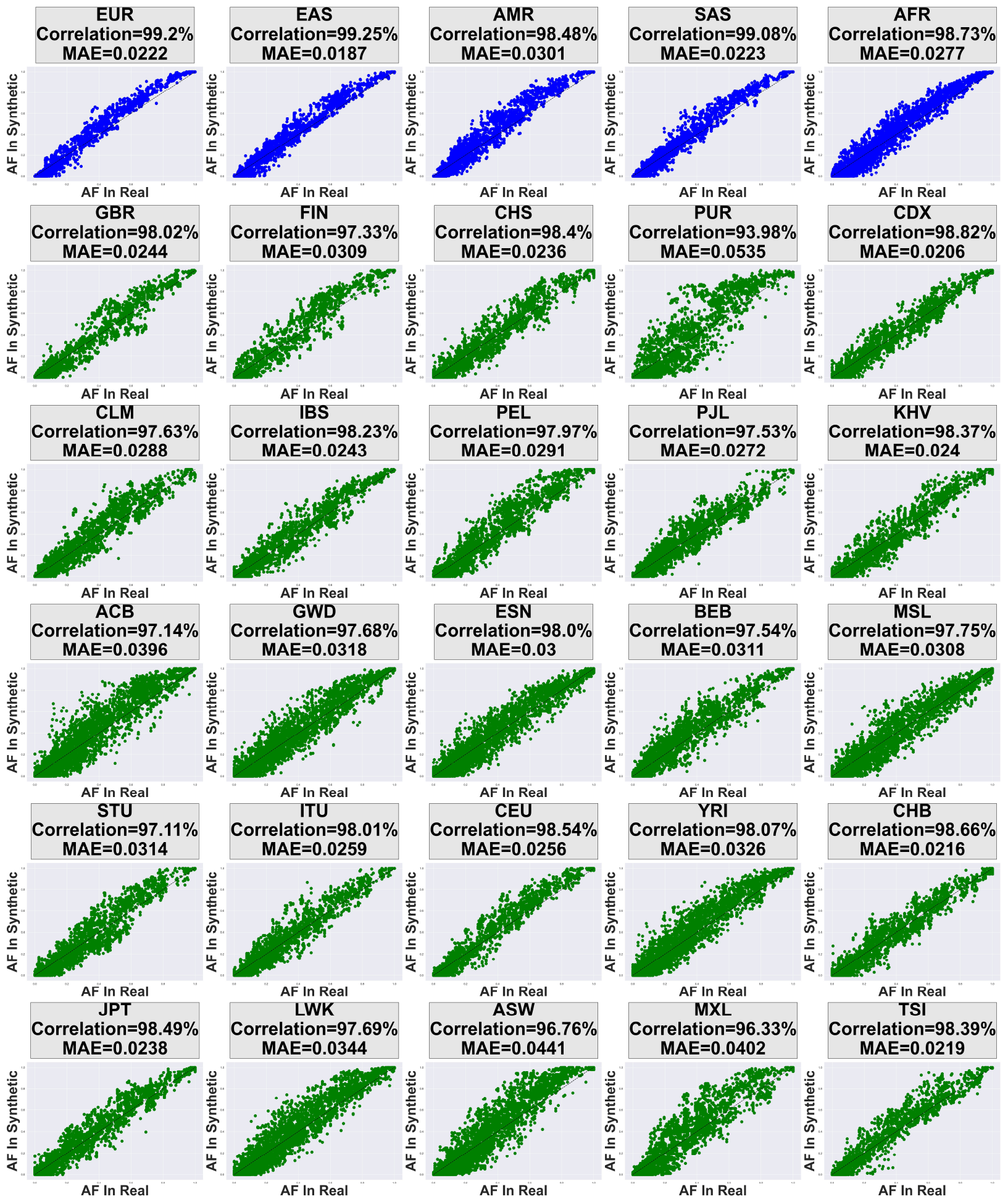
Population-specific allele frequencies in real and synthetic genomes. Allele frequencies were measured in each population for each of the 10, 000 SNPs for AGs and plotted against allele frequencies measured in real genomes (test and training set combined). The layout of each plot is similar to that used in Figure 9. The top five panels (blue) show a comparison of AGs generated by AC-GAN-Con, measuring allele frequencies in each of the five continental groups. The remaining 25 panels (green) show AGs generated by AC-GAN-Nat, measuring allele frequencies in 25 national populations. The GIH population from South Asia (Gujarati Indians in Houston) was omitted from this visualization to obtain a 5 *×* 5 layout. It produced very similar allele frequencies in AGs and real genomes as ITU (Indian Telugu in the U.K.).

Population-specific allele frequencies are particularly important when studying population differentiation, which is a fundamental component of population genetic analysis. Differentiation between two populations is represented in our setting through the difference between allele frequencies observed in the two populations across all 10, 000 SNPs. Thus, to examine how well our AC-GANs model differentiation, allele frequency differences were recorded in the real genomes using the training and test set combined, and then compared to allele frequency differences measured using AGs generated by our AC-GANs. Such comparisons are presented in Figure 11 for five pairs of continental groups and five pairs of national populations with varying degrees of differentiation. First, we see that AC-GAN-Con models allele frequency differences quite well, even between continents with lower genetic differentiation, such as America and East or South Asia (two upper left panels in Figure 11). Similarly, AC-GAN-Nat models allele frequency differences quite well between national populations from different continents (three lower right panels in Figure 11). We note that the accuracy in allele frequency difference is only mildly reduced relative to that obtained by AC-GAN-Con, despite the substantial reduction in the number of training samples per class. Not surprisingly, differentiation is much more difficult to model between national populations from the same continent because differentiation between such populations is typically more subtle. Nonetheless, AC-GAN-Nat was able to model allele frequency differences reasonably well even for several weakly differentiated populations within the same continent (two lower left panels in Figure 11). Importantly, the slight bias observed in allele frequencies (Figure 9) does not manifest itself in allele frequency differences, and as a result population differentiation is not systematically increased or decreased by the Genome-AC-GAN.

**Figure 11:**
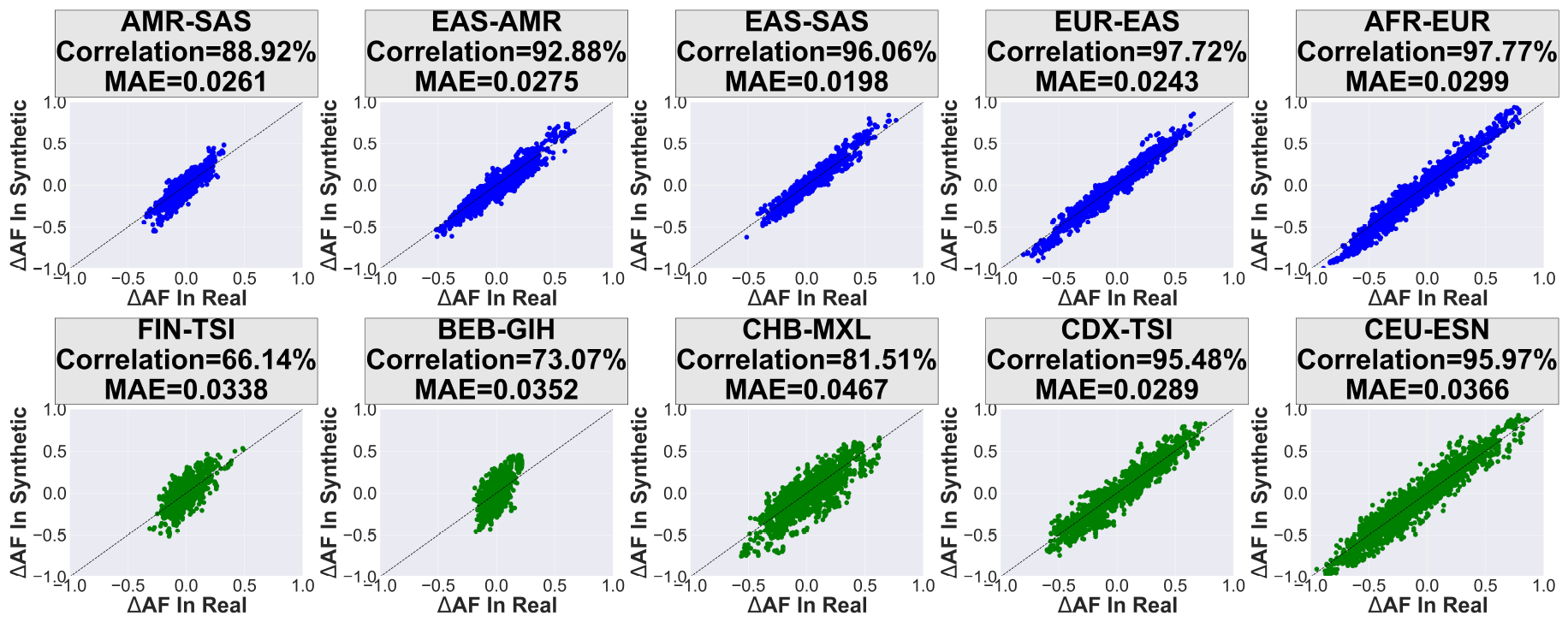
Population differentiation in real and synthetic genomes. Allele frequency differences were measured for five pairs of continental groups and five pairs of national populations in each of the 10, 000 SNPs. The setup and plot layout are similar to those in Figure 10, but instead of allele frequencies, allele frequency differences are plotted (in the range [−1, 1]). Measurements for real genomes are based on all 5, 006 genomes in the test and training set combined. The top five panels (blue) show a comparison of AGs generated by AC-GAN-Con for five pairs of continental groups, and the bottom five panels (green) show measurements obtained using AGs generated by AC-GAN-Nat for five pairs of national populations. Pairs were selected to cover different levels of population differentiation (As indicated by the range of allele frequency difference observed for each pair).

#### Linkage disequilibrium

Another key aspect of genetic variation is the correlation between alleles in different genomic loci caused by the way alleles are passed by inheritance from one generation to the next. Alleles in nearby loci are typically passed to the next generation together, unless they are unlinked by genetic recombination. This causes linkage disequilibrium (LD) between alleles in different loci, which has well-studied patterns along the genome [29]. In essence, LD measures the correlation between two loci, which tends to decay with distance. To compare patterns of LD in AGs to those observed in real genomes, all 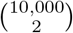 pairs of SNPs were partitioned into 50 bins according to the distance between them along the genome (from 1bp to ∼ 6Mb). Then, for every bin, an average LD was computed over all SNP-pairs in that bin for the real genomes in the test set, the real genomes in the training set, and the AGs produced by the five models in Table 2. The resulting patterns of LD are shown in Figure 12, and the performance of each model is summarized using the root mean square error (RMSE) from its vector of LD measurements and the one computed for the test set. As expected, the real genomes in the training set produced nearly identical patterns as observed in the test set (RMSE= 0.002). Furthermore, as observed in [40, 39], generative models produce the expected pattern of LD decay, but with consistently lower values of LD when compared to real genomes. Notably, the two AC-GANs produce higher average LD than any of the other three generative models across all bins. AC-GAN-Con performed particularly well in this respect, with RMSE= 0.032, which is roughly half of that measured for the previous best method (GAN19 with RMSE=0.063). Thus, the use of an AC-GAN appears to significantly improve the accuracy of LD in AGs, with the coarse-grained class labels of AC-GAN-Con performing better than the higher resolution labels of AC-GAN-Nat.

**Figure 12:**
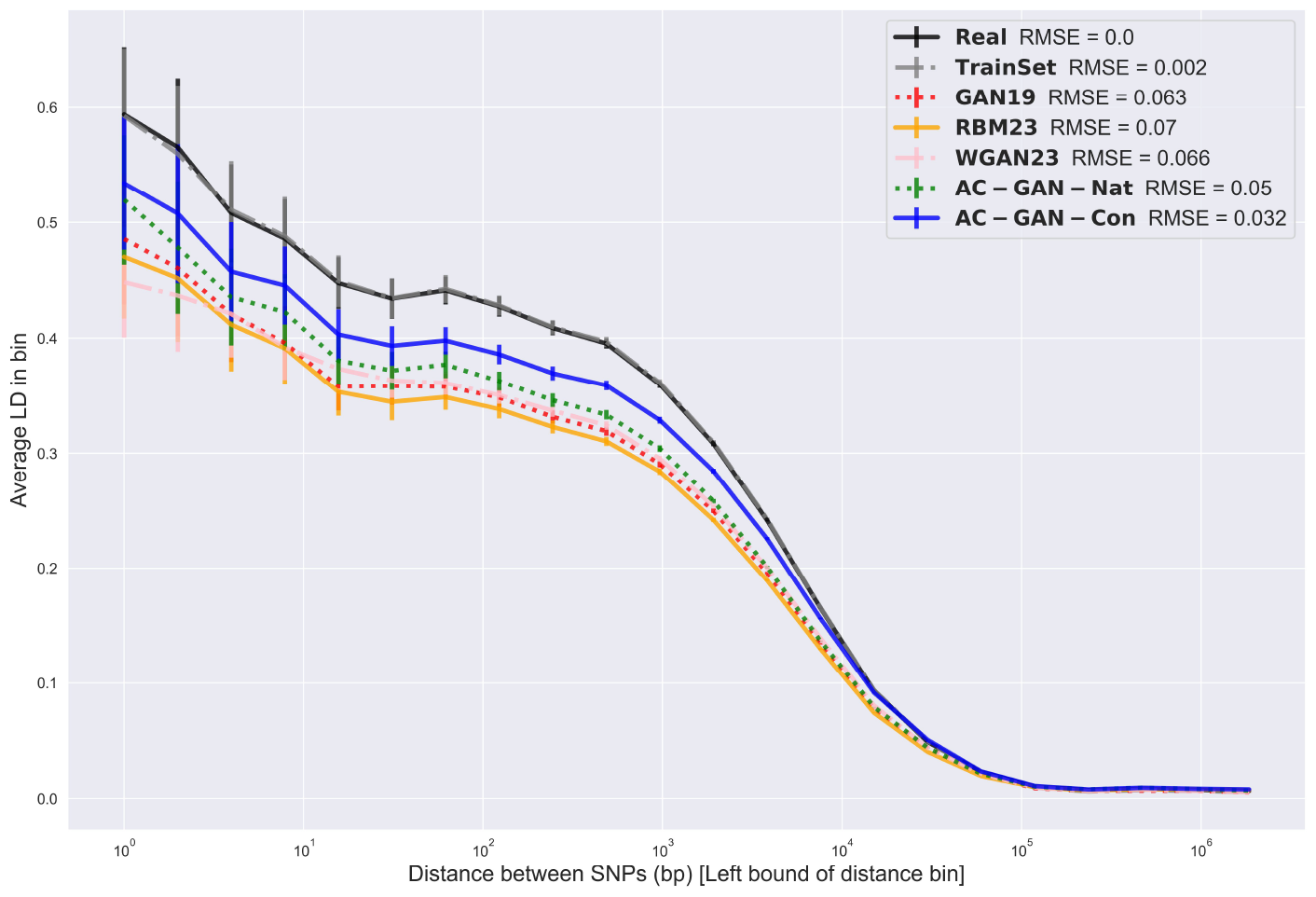
Linkage disequilibrium (LD) in AGs. All 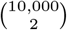 pairs of SNPs were partitioned into 50 bins according to the distance between them along the genome. Then, for every bin and every set of genomes (test set, training set, and five sets of AGs; see legend), the average LD and standard deviation were computed over all SNP-pairs in that bin. The plot shows average LD (Y-axis) as a function of distance between SNPs (X-axis in log-scale), with vertical bars representing standard deviations. The performance of each model was summarized using the root mean square error (RMSE) from its vector of LD measurements and the one computed for the test set.

### 4.3 Using synthetic genomes to augment training sets of classifiers

Finally, we wished to assess the potential advantages of utilizing synthetic genomes generated by the Genome-AC-GAN to improve classification. The premise of this experiment, as demonstrated in [38] for detecting COVID in chest X-rays, was that classification methods are often trained using a limited number of samples from each class, and synthetic samples generated by an AC-GAN can be used to boost the training signal. In our experiments, we considered the task of classifying genomes of individuals of African descent (AFR) into national populations. We chose this specific classification task, since we expected it to be relatively challenging due to lack of genetic differentiation between the seven African populations in the 1000 Genomes data set, as suggested by their 3D PCA (Figure 13). For this purpose, we re-trained a version of the Genome-AC-GAN only on AFR genomes, using *C* = 7 national populations as class labels. We employed the same training procedure described previously for AC-GAN-Con and AC-GAN-Nat. In particular, 80% of the AFR genomes were used for training (1, 056), and 20% were set aside for testing (266), maintaining this ratio across national populations. Training used the PLCE loss function with *ϵ* = 0.2, *α* = 0.1 and proceeded for 10, 000 epochs. The trained Genome-AC-GAN was then used to generate 1, 056 AGs (matching the size of the training set) that were later used to augment the training set in the classification task.

**Figure 13:**
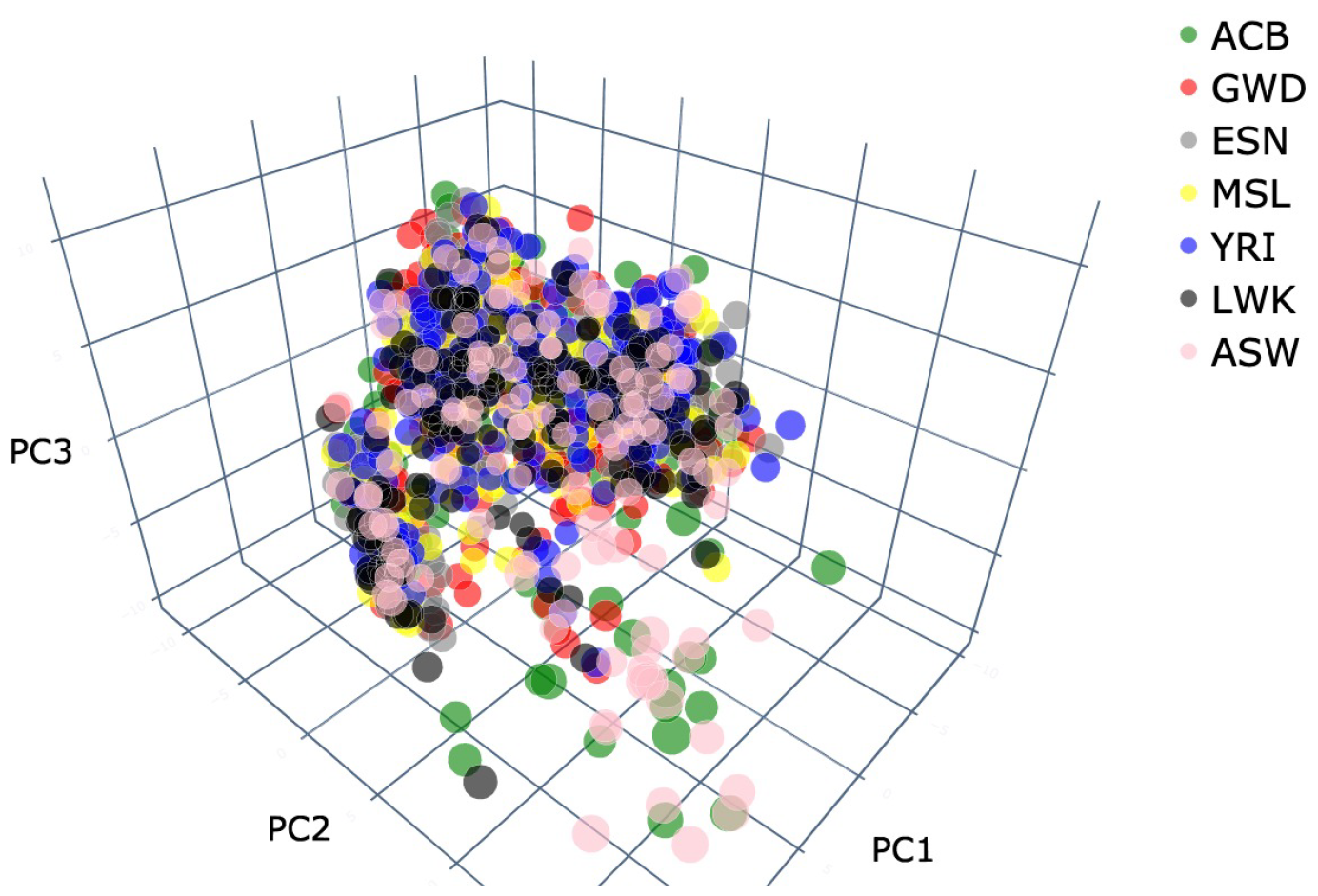
PCA of individuals of Afican descent. The 1, 322 genomes in the 1000 Genomes dataset are associated with individuals of African descent (AFR; see Table 1) were projected onto the three main PCs. Genomes are labeled in color according to their national population (see legend). The distributions of the seven national populations in this PCA largely overlap, indicating that the classification of these genomes into national populations is expected to be challenging.

We considered three fundamental models for classifying genomes into populations: a support vector classifier (SVC) [2] with a radial basis function kernel, a K-nearest neighbors (KNN) model [2] with *K* = 5, and a six-layer neural network (NN) [30]. The SVC and KNN were implemented using the appropriate functions in `scikit-learn` with default parameters. The NN consisted of six fully-connected hidden layers with the LeakyReLU activation function and *L*_2_ regularization, and an output layer with softmax transformation. It was trained using a PLCE loss function and the RMSprop optimizer, with batch size set to 512. These three classifiers were trained on various augmented versions of the training set, created by adding AGs generated by the AC-GAN in ten increments of 10% of the original training set size. Thus, the smallest training set had 1, 056 real genomes and no AGs, and the largest training set had 1, 056 real genomes and 1, 056 AGs. In each case, AGs were resampled for each of the seven national populations from the collection of 1, 056 AGs according to their prevalence in the original training set (Table 1). For each level of augmentation (0%–100%), we conducted 50 different replicate experiments to capture the randomness of the classification methods and resampling of AGs. Throughout all experiments, the exact same training procedure was maintained for every classifier, to ensure consistency of the results. Finally, the accuracy of the three trained classifiers was evaluated in each replicate using the 266 genomes in the test set.

First, we observe a consistent pattern of improvement in classification accuracy for all classifiers as more AGs are added to the training set (Figure 14). This is most prominent for the KNN model, which improves from an average accuracy of 28.9% on the original training set to an average accuracy of 34.2% on the training set doubled by adding AGs. The classification accuracy of the SVC model also increases consistently as more AGs are added (from 32.3% to 36.1%). The NN model shows somewhat different behavior, with classification accuracy increasing from 31.4% to 36.5% when adding 30% AGs, but then maintaining roughly the same level of accuracy when more AGs are added. We then examined the improvement in classification accuracy for each national population when doubling the training set by adding AGs (Figure 15). The three classifiers show increased accuracy when trained on augmented data for all seven populations, with the exception of the classification of the MSL population by the KNN model. Interestingly, the classification accuracy of the same population is more than doubled for the NN model (from 12.8% to 27.4%). In 13 out of the total 21 cases, we observed a relative increase in accuracy larger than 20%, indicating the utility of augmentation particularly in classification tasks characterized by initially low accuracy rates. While this experiment illustrates the utility of synthetic augmentations in specific models and scenarios, it also indicates that the outcomes and required level of augmentation are variable for different classifiers.

**Figure 14:**
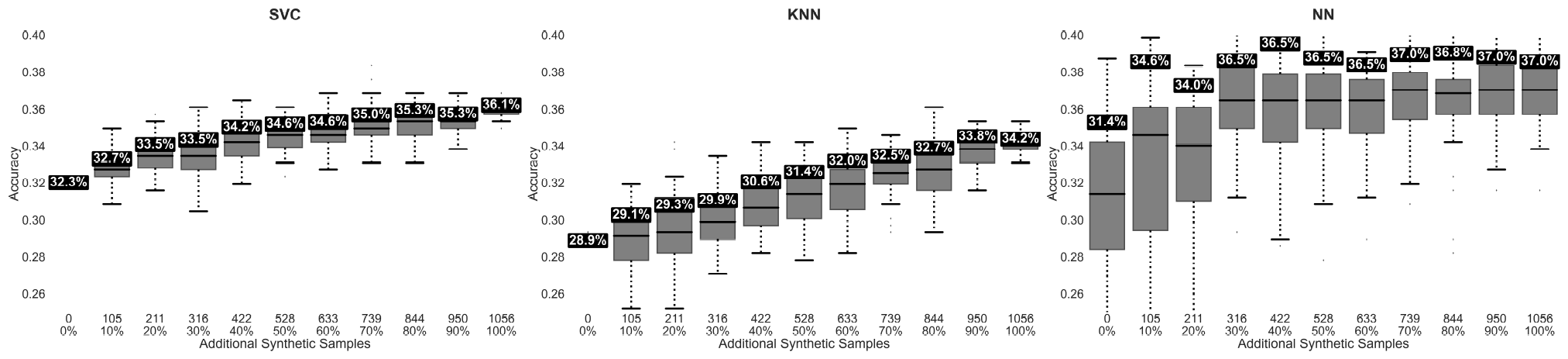
Classification accuracy as a function of number of AGs added to the training set. The classification accuracy of three models (SVC, KNN, and NN) is evaluated as a function of the number of AGs added to the training set. Each box plot depicts the distribution of classification accuracy across 50 replicates. The middle bar in each box plot signifies the median, while the box itself spans the interquartile range (IQR), encapsulating the central 50% of the data, and the whiskers extend to the minimum and maximum values. The mean accuracy is specified at the top of each box plot, and the number of AGs added to the training set is specified at the bottom together with and their percentage from the original set of 1, 056 real genomes.

**Figure 15:**
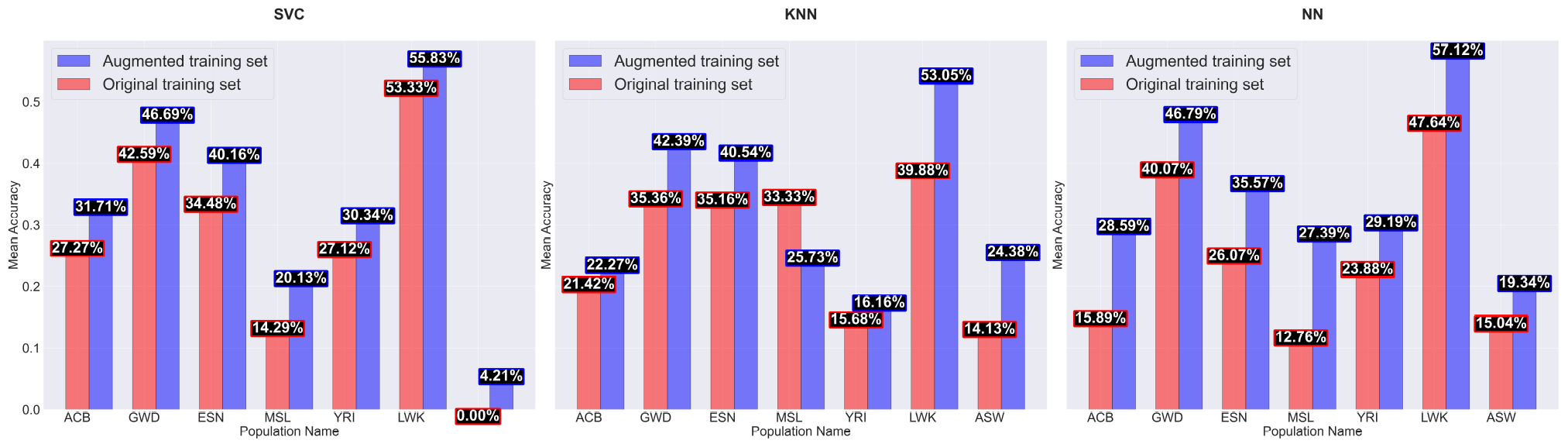
Population-specific classification accuracy when augmenting the training set with 100% AGs. The classification accuracy of three models (SVC, KNN, and NN) is evaluated for each of the seven national populations (which are the class labels) when trained on the original training set (red) and a training set doubled by adding AGs generated by the Genome-AC-GAN (blue). Each bar represents the mean classification accuracy across 50 replicate experiments, as measured on the test set (see Figure 14)

## 5 Conclusion and discussion

This paper introduces the Genome-AC-GAN model, a novel approach for generating artificial genomes. Unlike previous generative models, the AC-GAN approach allows Genome-AC-GAN to explicitly model sub-populations within a larger cohort. The utility of an AC-GAN depends on the ability to effectively train the discriminator to classify samples into these sub-populations. Careful finetuning of the polyloss cross entropy loss function allowed us to improve the accuracy of classification (Figure 3), and as a result improve the realism of the generated genomes. Our experimental analysis showcases the capabilities of the Genome-AC-GAN in generating artificial genomes closely resembling real genomes, typically slightly outperforming other recently published models. The only noteable exception to this rule was that the WGAN model of [39] produces slightly more accurate allele frequencies than Genome-AC-GAN (Figure 9). On the other hand, Genome-AC-GAN produced much more realistic LD patterns than the WGAN or any other model (Figure 12). This is an important (and somewhat unexpected) advantage of the Genome-AC-GAN, since this was highlighted as one of the main deficiencies of previous generative models for genomic data [40].

The fact that the Genome-AC-GAN considers sub-populations within a larger cohort enables it to model more complex distributions of genetic variation, and likely contributes to its improved accuracy compared to other methods. It also provides the generative model with the unique capability of modeling genetic differences between populations (Figures 7 and 11). While a standard generative model might sacrifice accuracy in smaller populations to improve overall accuracy, the AC-GAN approach can be used to guide the model toward more accurate modeling of under-represented populations. This was apparent in our results for AC-GAN-Nat, in which some populations had fewer than 200 training samples. An important consideration when implementing an AC-GAN is to decide on the number of classes and their resolution. Higher-resolution classes can potentially allow more complex modeling, but result in fewer training samples per class. Comparing our two Genome-AC-GANs, we see that the higher-resolution version (AC-GAN-Nat) was slightly more accurate in modeling overall genetic variation and allele frequencies (Figures 5 and 9), but the lower-resolution version (AC-GAN-Con) more accurately modeled LD (Figure 12). That said, both versions performed well in all measurements, confirming the robustness of the AC-GAN approach to the number of classes and their size.

There are several potential applications of AC-GANs in genomics. Here, we examined the potential use in augmenting training sets for classification tasks (Figures 14 and 15). We showed that data augmentation can be used to improve classification accuracy in challenging tasks that involve similar classes with a small number of training samples. While the improvement was not dramatic (e.g., from 32% to 36% for SVC), it was consistent and robust across different classifiers and different classes. It is important to note that this approach has an inherent limitation, since the artificial genomes used in augmentation do not carry information that is not present in the original training set. Our results suggest that the augmented genomes effectively model this information in a way that improves the classifiers’ capabilities to process it. One potential application where this approach might be useful is the study of genetic contributors to rare phenotypes. While standard approaches require a large number of sequenced genomes [8, 41], by utilizing an AC-GAN that considers phenotypes as classes, it might be possible to make better use of smaller numbers of genomes. More generally, developing generative methods that jointly model genotypes and phenotypes remains a major challenge that can be addressed using AC-GANs.

